# A lymphoid tissue chemokine checkpoint prevents loss of CD8^+^ T cell functionality

**DOI:** 10.1101/2024.09.19.613830

**Authors:** Lukas M. Altenburger, Daniela Claudino Carvoeiro, Philippe Dehio, Jianwen Zhou, Chiara Laura, Mitali Katoch, Caroline Krüger, Juliana Barreto de Albuquerque, Petra Pfenninger, Jose Martínez Magdaleno, Jun Abe, Matthias Mehling, Jörn Dengjel, Matteo Iannacone, Ali Hashemi Gheinani, Jens V. Stein

## Abstract

The generation of effector CD8^+^ T cells (T_EFF_) requires activation of naive CD8^+^ T cells (T_N_) by dendritic cells (DCs) within lymphoid tissue. To date, it remains elusive how the duration of T_N_-DC interactions and integration of activation signals are controlled *in vivo*. Here, we report that lymphoid stroma-secreted ligands for CCR7 constrained interaction duration by gradually inducing CD8^+^ T cell release from DCs. At late time points of interactions, CCR7 ligands repositioned the F-actin-promoting factor DOCK2 away from the DC interface to enable CD8^+^ T cell detachment, proliferation onset and acquisition of cytotoxicity. Lack of CCR7 signaling, as during *ex vivo* activation or in chronically inflamed lymphoid tissue, caused sustained T cell-DC interactions, and generated dysfunctional T_EFF_ with high expression of inhibitory receptors, impaired antimicrobial activity, and poor recall responses. In sum, our findings uncover that lymphoid stromal chemokines act as built-in “disruptors” of T cell-DC interactions for long-term preservation of T_EFF_ functionality.

## Introduction

During infections with intracellular pathogens, the adaptive immune system efficiently expands rare Ag-specific clones from a pool of naive CD8^+^ T cells (T_N_) containing millions of distinct T cell receptor (TCR) specificities (*1*, *2*). The chemokine receptor CCR7 is expressed on T_N_ and activated dendritic cells (DCs) and constitutes a critical mediator of rapid clonal selection. CCR7 directs the colocalization of both cell types in the paracortex of lymph nodes (LNs) and other secondary lymphoid organs (SLOs) (*3*), where ICAM-1^+^ fibroblastic reticular cells (FRCs) secrete its ligands CCL19 and CCL21 (*4–6*). Within the paracortex, CCR7 and the ICAM-1 receptor LFA-1 coordinate dynamic T_N_ scanning of peptide-major histocompatibility complexes (pMHC) presented on DCs (*7–9*). Engagement of the TCR and CD28 by cognate pMHC and CD80/CD86, respectively, presented on activated DCs leads to T_N_ arrest and the formation of an immunological synapse (IS). During their interactions, DCs instruct a transcriptional program in CD8^+^ T cells that licenses their differentiation to cytotoxic effector T cells (T_EFF_). The duration of their interactions with DCs determines the level of activation signal integration from engaged TCR and costimulatory receptors, thereby decisively shaping T_EFF_ differentiation (*10*, *11*). Thus, whereas a short TCR stimulus suffices to induce a CD8^+^ T cell division program without further stimulation (*12*, *13*), prolonged TCR stimulation increases the magnitude of CD8^+^ T cell expansion (*14–16*). However, chronic TCR signaling leads to dysfunctional T cell activation with phenotypic features of T cell exhaustion, characterized by reduced proliferation, expression of inhibitory receptors such as programmed cell death protein-1 (PD-1), and impaired effector function (*17*, *18*). Despite the fundamental impact of activation signal integration on T_EFF_ differentiation (*19*, *20*), there is remarkably little information on how T_N_-DC interactions are timed within lymphoid tissue.

## Results

To investigate the mechanism underlying interaction duration, we compared *in vitro* and *in vivo* models of T cell activation (**Figure 1A**). We used the Nur77^GFP^ reporter mouse strain, where GFP expression reports recent antigen receptor signaling activity (*21*), and backcrossed these mice to *Discosoma* red fluorescent protein (DsRed)^+^ OT-I TCR tg T cells, which recognize the OVA_257-264_ (SIINFEKL) peptide in the context of H-2K^b^ (*22*). For *in vivo* priming, we adoptively transferred cell tracker violet (CTV)-labeled Nur77^GFP^ DsRed^+^ OT-I T cells into C57BL/6 recipients that contained subcutaneously (s.c.) injected DCs pulsed with 100 nM OVA_257-264_. Previous studies have shown that this pMHC level rapidly induces stable interactions, followed by gradual detachment over the next 18-32 h (*23*). In accordance, we found that GFP levels rapidly decreased across daughter cell generations at 48-72 h post T cell transfer (**Figures 1B and 1C; Figure S1A**). This finding is consistent with CD8^+^ T cell release from DCs prior to the initiation of cell division (*23–25*), suggesting only limited engagement of proliferating T cells with DCs. In contrast, GFP levels decreased much less in daughter cell generations of Nur77^GFP^ OT-I T cells activated *in vitro* with 100 nM OVA_257-264_-pulsed DCs, suggesting that outside the lymphoid microenvironment, CD8^+^ T cells continue TCR signaling as they proliferate (**Figures 1B and 1C**). The GFP MFI decay kinetics of *in vitro* activated Nur77^GFP^ DsRed^+^ OT-I T cells closely resembled those of continuously stimulated CD8^+^ T cells using mAbs against CD3ε and the costimulatory receptor CD28 (**Figures 1B and 1C**). Continuous *in vitro* activation also led to a lower number of cell divisions than *in vivo* stimulation at 48 h (**Figure 1D**) but did not correlate with decreased TCR surface levels (**Figure S1B**). Consistent with the flow cytometry results, intravital imaging of the LN paracortex identified only few transient T cell-DC interactions at 22-28 h post T cell transfer, whereas a second cohort of T_N_ added at this time engaged in stable interactions with the same DCs (**Figure 1E; Video S1**). In contrast, *in vitro* divided CFSE^low^ OT-I daughter cells remained attached to OVA_257-264_-pulsed DCs at 48 h of co-culture (**Figure 1E**). In support of restricted T cell-DC interactions in SLOs, we observed a rapid decay of Nur77^GFP^ signal in dividing OT-I T cells isolated from LNs and spleens at 40-48 h post infection (p.i.) with lymphocytic choriomeningitis virus-OVA (LCMV-OVA) and *Listeria monocytogenes*-OVA (Lm-OVA), respectively (**Figures 1F and 1G**). Taken together, *in vitro* activated CD8^+^ T cells continue to interact with DCs during proliferation, resulting in prolonged TCR signaling throughout cell division cycles. In contrast, the lymphoid tissue microenvironment contains checkpoints that promote CD8^+^ T cell detachment from DCs before proliferation onset (*11*).

**Figure 1.**
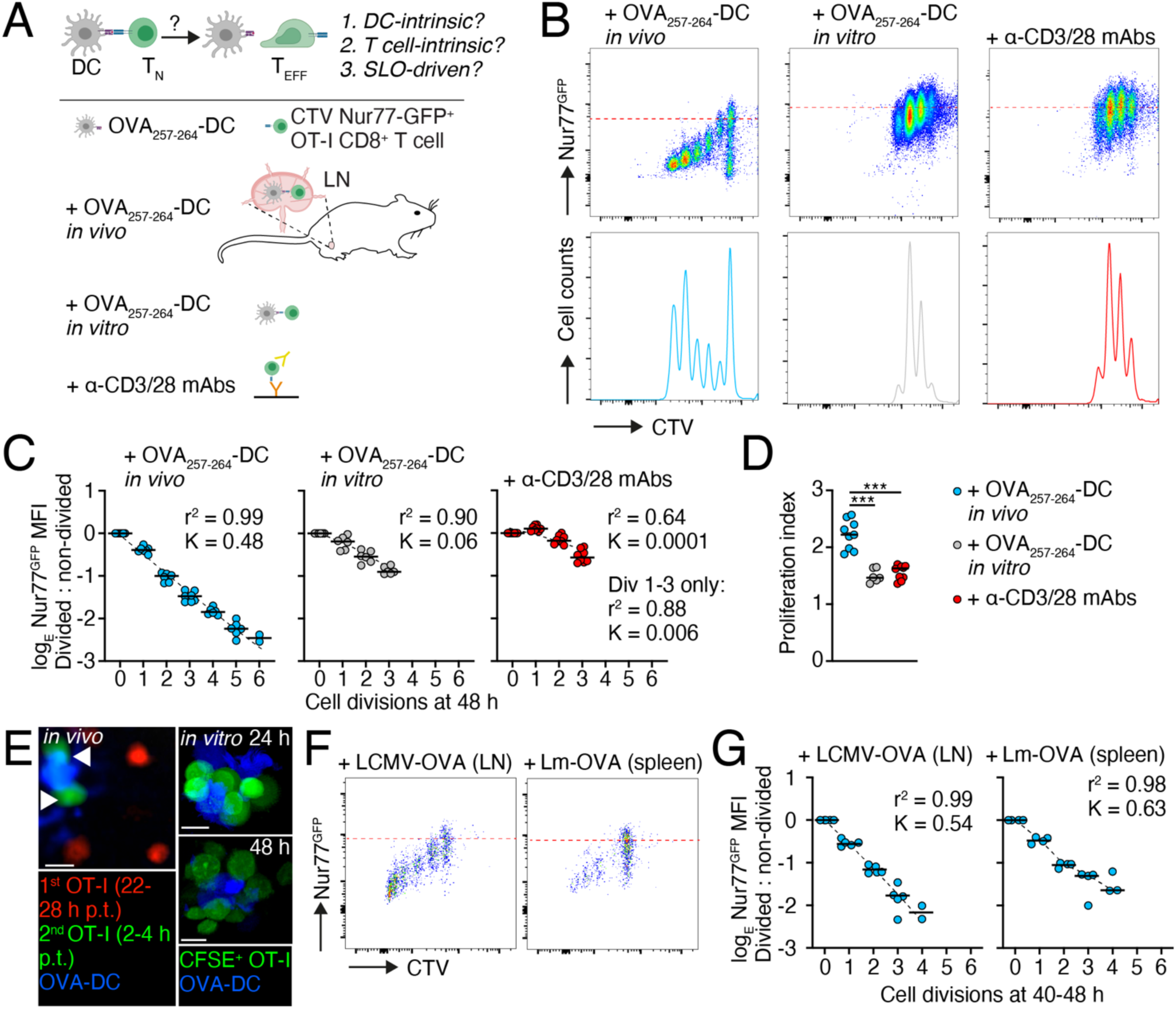
T cell-DC interactions are capped within lymphoid tissue. **A.** Experimental layout. **B.** Flow cytometry of CTV-labeled Nur77^GFP^ OT-I T cells at 48 h post activation. **C.** GFP MFI decay of activated Nur77^GFP^ OT-I T cells at 48 h post activation. **D.** Proliferation index of Nur77^GFP^ OT-I T cells at 48 h post activation. **E.** Representative intravital image of OT-I T cells transferred 22-28 h (1^st^) or 2-4 h (2^nd^) before LN imaging (p.t., post transfer), and confocal images of *in vitro*-activated OT-I T cells interacting OVA_257-264_-pulsed DCs at 24 and 48 h post activation. Arrowheads depict 2^nd^ cohort OT-I T cells attached to DCs. Scale bars, 10 µm. **F.** Flow cytometry of CTV-labeled Nur77^GFP^ OT-I T cells isolated from LNs and spleens at 40-48 h p.i. with LCMV-OVA and Lm-OVA, respectively. **G.** GFP MFI decay of activated Nur77^GFP^ OT-I T cells at 40-48 h post *in vivo* activation. Red dotted lines in B and F depict GFP expression in activated, non-divided OT-I T cells. K, rate constant; Div, division. Data in C, D and G are pooled from at least two independent experiments and analyzed using ANOVA with Tukey’s post-test (D). ***, p < 0.001.

### CCR7 ligands limit T cell-DC interaction duration

While chemokines, in particular CCR7 ligands, are abundantly expressed in the LN paracortex to promote T cell motility, their role as putative regulators of CD8^+^ T cell-DC interaction duration has not been addressed to date. We therefore co-incubated OT-I T cell and OVA_257-264_-pulsed DCs for 4, 10 or 24 h before exposing them to gradients of the CCR7 ligand CCL21 in Transwell assays. To restrict the chemokine-induced migratory response to CD8^+^ T cells, we used CCR7^-/-^ DCs as APCs (**Figure 2A**). At 4 h after DC engagement, OT-I T cells completely lost their migratory response to CCL21, despite retaining high surface CCR7 levels (**Figures 2B and 2C**). At 10 and 24 h after DC engagement, OT-I T cells gradually recovered migration towards CCL21, although CCR7 surface levels slightly decreased over this period (**Figures 2B and 2C**). Migrated OT-I T cells remained undivided and expressed the early activation marker CD69, indicative of TCR engagement (**Figure 2D**). We observed a similar pattern of transient unresponsiveness towards CCL21 in P14 TCR tg CD8^+^ T cells interacting with LCMV gp_33-41_-pulsed DCs (**Figure S2A**) and when using CCR7^+/+^ DCs as APCs (**Figure S2B**). OT-I T cells activated by DCs presenting a low affinity peptide-MHC (SIIQFEKL peptide, “Q4”) (*26*) also displayed a transient lack of CCR7-mediated chemotaxis (**Figure S2C**). To investigate a potential role of DCs in controlling detachment kinetics, we added a fresh cohort of OT-I T_N_ to a 20 h co-culture of OVA_257-264_-pulsed DCs and OT-I T cells and exposed the mixed cultures 4 h later to a CCL21 gradient in a Transwell assay. While OT-I T cells that had interacted for 24 h (cohort 1) had recovered CCL21 responsiveness, the second cohort of OT-I T cells (4 h of interactions) exhibited severely reduced migration (**Figure S2D**).

**Figure 2.**
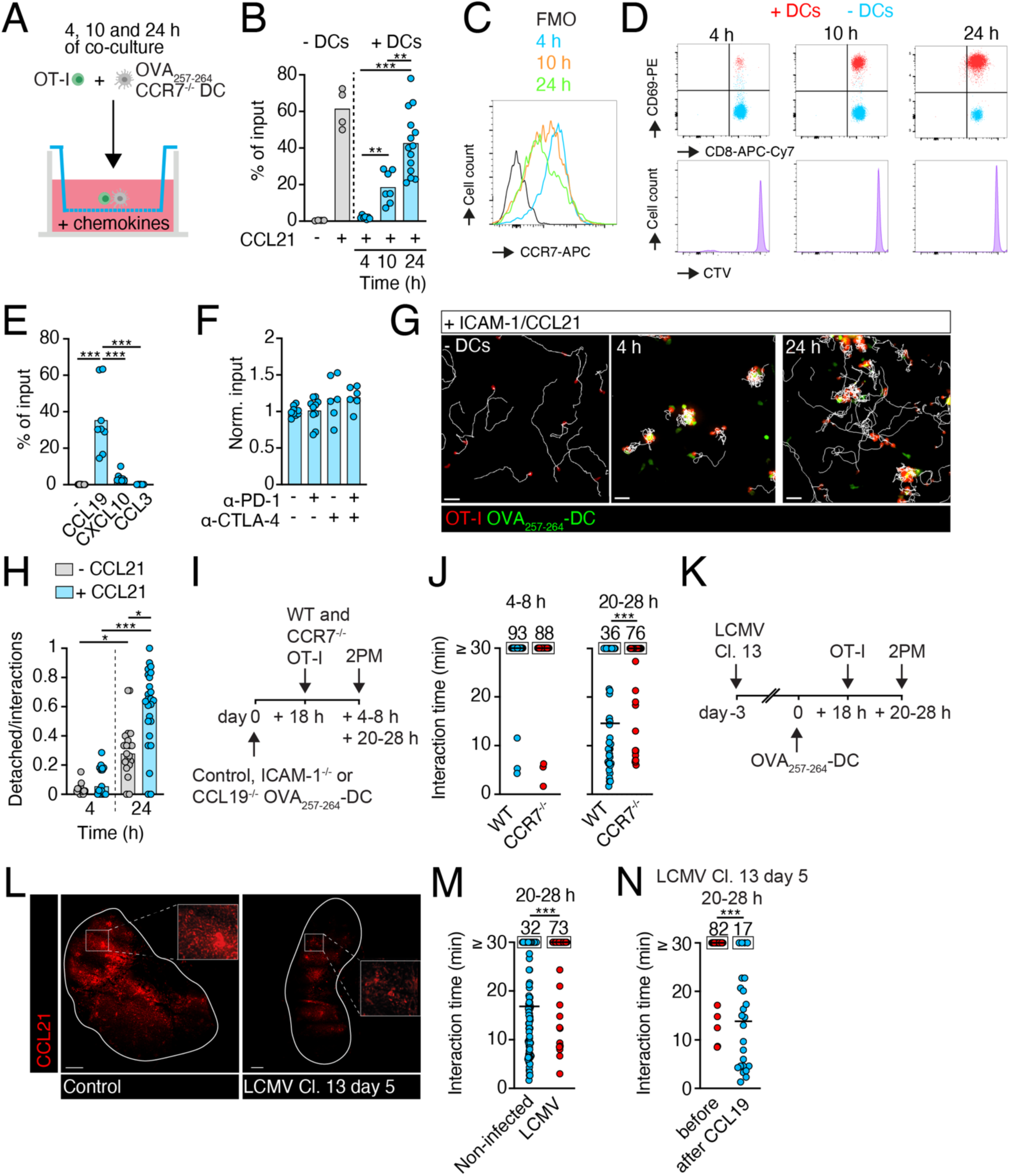
CCR7 ligands disrupt late CD8^+^ T cell-DC interactions. **A.** Experimental layout of T cell-DC co-culture. **B.** Chemotaxis of OT-I T cells left unstimulated or interacting for 4, 10 or 24 h with OVA_257-264_-pulsed DCs towards CCL21. **C.** CCR7 expression on OT-I T cells interacting with OVA_257-264_-pulsed DCs. **D.** CD69 expression of CTV-loaded OT-I T cells that have migrated to CCL21 with or without DCs in top chamber. **E.** Chemotaxis of OT-I T cells interacting for 24 h with OVA_257-264_-pulsed DCs towards CCL19, CXCL10 (ligand for CXCR3) and CCL3 (ligand for CCR5). **F.** Normalized chemotaxis of OT-I T cells interacting for 24 h with OVA_257-264_-pulsed DCs towards CCL21 in presence of blocking mAbs against PD-1 and/or CTLA-4. **G.** Exemplary tracks of unstimulated OT-I T cells or OT-I T cells interacting for 4 or 24 h with OVA_257-264_-pulsed DCs on CCL21 + ICAM-1-coated plates under agarose. Scale bar, 30 µm. **H.** Detachment events for OT-I T cells normalized to all OT-I T cells in a field of view. **I.** Experimental layout of intravital imaging (2PM). **J.** Interaction times of control and CCR7^-/-^ OT-I T cells with OVA_257-264_-pulsed DCs *in vivo*. **K.** Experimental layout of intravital imaging in LCMV Clone 13-infected LNs. **L.** CCL21 staining of control and LCMV-infected LN sections. Inserts depict paracortical CCL21 signal. Scale bar, 300 µm. **M.** Interaction times of OT-I T cells with OVA_257-264_-pulsed DCs in control and infected LNs. **N.** Interaction times of OT-I T cells with OVA_257-264_-pulsed DCs in infected LNs before and after CCL19 injection. Numbers in J, M, and N indicate percentage of long-lasting (≥ 30 min) interactions. Data are pooled from at least two independent experiments and analyzed using ANOVA with Dunnett’s post-test (B, E, F, H) and Kruskal-Wallis with Dunn’s post-test (J, M, N). *, p < 0.05; **, p < 0.01; ***, p < 0.001.

Given the overlapping yet distinct signaling cascades triggered by CCL19 versus CCL21 (*27*), we assessed the detachment potential of CCL19. In parallel, we tested whether CXCR3 and CCR5 ligands enforce T cell detachment from DCs, since CD8^+^ T cells increase expression of these chemokine receptors in reactive LNs (*28*, *29*). While CCL19 promoted OT-I T cell migration after 24 h of DC co-culture similarly to CCL21, ligands for CXCR3 and CCR5 did not induce T cell chemotaxis (**Figure 2E**). In accordance, *in vivo* activated, undivided OT-I T cells were CCR7^high^ CXCR3^low^, and increased CXCR3 levels only after cells had divided at least two times (**Figure S2E**). Finally, we addressed whether the immune checkpoints PD-1 and CTLA-4 might contribute to early OT-I T cell release from DCs. mAb blockade of PD-1 alone or in combination with anti-CTLA-4 mAb did not affect CCL21-induced OT-I T cell detachment at 24 h post initial contact (**Figure 2F**). In agreement, PD-1 and CTLA-4 levels remained low on undivided cells during *in vivo* activation (**Figure S2F**; not shown). These data suggest only a limited role for these immune checkpoints in restricting the initial CD8^+^ T cell priming in lymphoid tissue, in contrast to their well-described function in reducing T_EFF_-APC interactions in the late effector phase (*30*).

T cell migration on CCL19- and CCL21-secreting ICAM-1^+^ FRCs within the LN paracortex is best described as a guided random walk (*31–33*), where, unlike in the Transwell system, CCR7 ligands do not form a gradient (*34*). We therefore assessed how CD8^+^ T_N_-DC interaction duration was affected in an *in vitro* under-agarose assay on plates, which were uniformly coated with ICAM-1 and CCL21. While T_N_ showed high motility on CCL21 and ICAM-1-coated substrates in the absence of DCs, they rapidly stopped and continuously interacted with OVA_257-264_-pulsed CCR7^-/-^ DCs at 4 h, irrespective of the presence or absence of CCL21 (**Figures 2G and 2H; Video S2**). In contrast, after 24 h of interactions, CCL21 significantly enhanced OT-I T cell detachment from DCs (**Figures 2G and 2H; Video S2**).

Besides chemokines, lymphoid tissue contains other factors that potentially interfere with T cell-DC contact stability, such as regulatory CD4^+^ T cells or anti-inflammatory stromal cells (*35*, *36*). To selectively examine the role for CCR7 ligands during *in vivo* T_N_-DC priming, we compared interactions of WT and CCR7^-/-^ OT-I T_N_ with OVA_257-264_-pulsed DCs using intravital imaging (**Figure 2I**). While lack of CCR7 decreases T_N_ homing and interstitial motility, we took advantage of the fact that adoptively transferring an excess of CCR7^-/-^ over WT T_N_ allows comparing both populations in the same field of view (*37*, *38*), and that a high density of pMHC^+^ DCs compensates for decreased T_N_ motility to permit robust engagement frequencies (*39*). Intravital imaging of reactive LNs at 2-8 h post T_N_ transfer confirmed that most WT and CCR7^-/-^ OT-I T_N_ were stably attached to DCs, defined by continuous interactions throughout the observation period of 30 min (**Figure 2J**). At 20-28 h post transfer, while only a minority of WT OT-I T cells remained stably engaged with DCs, most CCR7^-/-^ OT-I T cells continued in stable interactions with DCs (**Figure 2J**). We observed a comparable delay in CCR7^-/-^ OT-I cell detachment when using ICAM-1^-/-^ DCs as APCs, suggesting that CCR7 ligands caused the release of ICAM-1-independent contacts as efficiently as contacts with WT DCs (not shown).

As a complementary approach to manipulate *in vivo* chemokine signaling, we exploited the fact that microbial infections lead to a marked decrease in stromal CCL19 and CCL21 expression within lymphoid tissues at 5-8 days p.i. (*40*, *41*). Despite its prevalence across multiple acute and chronic infection models, as well as in tumor-draining LNs (*42*, *43*), the impact of loss of stromal CCR7 ligands on T cell priming has not been explored. To provide comparable Ag levels on DCs as in non-infected recipients, we s.c. injected 100 nM OVA_257-264_-pulsed DCs into footpads of mice on day 3 after systemic infection with LCMV clone 13 (Cl. 13), followed by OT-I T cell transfer 18 h later (**Figure 2K**). At 20-28 h post OT-I T cell transfer (= day 5 p.i.), we performed intravital imaging of their interactions with OVA_257-264_-pulsed DCs, i.e., under conditions where the DCs are identical to control conditions but the environment is altered. At this time point, CCL21 levels were strongly reduced in the T cell zone of reactive LNs (**Figure 2L**). This finding correlated with significantly higher frequencies of stable OT-I T cell interactions to OVA_257-264_-pulsed DCs as compared to those seen in LNs in non-infected recipients (**Figure 2M**). Next, we locally replenished CCL19, which rapidly drains to sentinel LNs after s.c. administration (*37*). CCL19 administration quickly (< 1 h) restored T cell detachment from DCs in LNs of infected mice (**Figure 2N**). In sum, our data suggest that following an early stop signal by TCR engagement (*44*), CCR7 acts as a regulator of interaction duration as T cells gradually regain responsiveness to its lymphoid stroma-produced ligands, with similar kinetics of detachment under both *in vitro* and *in vivo* conditions (*23–25*, *45*, *46*). This process is disturbed in inflamed lymphoid tissue with low CCR7 ligand levels and could be an important driver of dysfunctional T cell activation during chronic infections and cancer.

### CCR7-induced repositioning of the Rac activator DOCK2 triggers T cell release from DCs

We asked how transient desensitization of T cells against CCR7 ligands during activation allowed for their initial stabilization of DC contacts. Pharmacological inhibition of G-protein receptor kinases (GRKs), which govern chemokine receptor internalization and lymphocyte migration (*47*), did not rescue T cell migration to CCL21 at 4 h post DC engagement (**Figure S3A**), consistent with continued surface expression of CCR7 at this time point (**Figure 2C**). Previous studies have provided evidence for a correlation between intracellular Ca^2+^ flux and lymphocyte arrest (*48–51*). We therefore examined Ca^2+^ levels in OT-I T cells interacting with OVA_257-264_-pulsed DCs in under agarose assays. OT-I T cells exhibited frequent Ca^2+^ pulses with high amplitudes at early but not late time points of interaction (**Figures S3B-D; Video S3**), and addition of the Ca^2+^ ionophore ionomycin decreased T_N_ migration to CCL21 in the absence of TCR engagement (**Figure S3E**).

Thus, early arrest correlates with TCR-driven Ca^2+^ flux but not chemokine receptor internalization. Furthermore, an analysis of published phosphoproteomic data sets (*52*) uncovered multiple TCR-induced changes in early and late phosphorylation patterns of proteins involved in cytoskeletal regulation, which might contribute to transient T cell arrest, including DOCK2, a Rac guanine exchange factor (GEF) expressed in hematopoietic cells (*53–55*) (**Figures S3F-I**). DOCK2 plays a dual role for both CCR7- and TCR-triggered F-actin polymerization (**Figure 3A**) (*39*, *53–57*). To further define the intracellular DOCK2 localization during T cell-DC interactions and release, we employed a microfluidic chamber in which, following either 4 or 24 h of co-culture, OVA_257-264_- pulsed CCR7^-/-^ DCs and OT-I T cells expressing a DOCK2-GFP fusion protein (*58*) were exposed to a CCL21 gradient (**Figure 3B**). We confirmed the lack of response to a CCL21 gradient in WT OT-I T cells interacting for 4 h with OVA_257-264_-pulsed DCs in this system, followed by an efficient detachment response after 24 h of interactions (**Figure 3C**). Using live cell imaging, we found that at 4 h post T cell engagement, DOCK2-GFP was sequestered at the interface with DCs even when cells were exposed to a CCL21 gradient, which correlated with few detachment events (8% of total interactions; **Figures 3D and 3E; Video S4**). At 24 h post engagement, when the frequency of DOCK2-GFP OT-I T cells detaching from DCs increased to 51%, we observed DOCK2 relocalization away from the DC interface immediately before T cell detachment in the majority (76%) of detachment events (**Figures 3D and 3E; Video S4**). To examine whether DOCK2 relocalization required CCL21 sensing, we deleted CCR7 on DOCK2-GFP OT-I T_N_ using CRISPR/Cas9. At 24 h of interactions, CCR7^del^ DOCK2-GFP OT-I T cells showed fewer detachment events from OVA_257-264_-pulsed DCs after exposure to a CCL21 gradient as compared to control cells, with only 25% of detachments preceded by DOCK2 relocalization away from the DC interface compared to 87% for control OT-I T cells (**Figure 3F**). Finally, we examined whether DOCK2 relocalization was required for T cell detachment from DCs. Microfluidic chamber imaging of DOCK2^-/-^ T cell-DC pairs revealed impaired detachment upon CCL21 exposure at 24 h post interactions (**Figures 3G and 3HI; Video S5**). In support of this, Transwell assays revealed a substantial reduction of CCL21-driven detachment of DOCK2^-/-^ OT-I T cells interacting with OVA_257-264_-pulsed DCs as compared to WT OT-I T cells (**Figures 3I and 3J**). We confirmed these findings using intravital imaging of LNs containing OVA_257-264_-pulsed DCs, as the percentage of stable T cell-DC interactions at 20-28 h post T cell transfer was increased from 49% in WT to 71% in DOCK2^-/-^ OT-I T cells (**Figure 3K**). Taken together, our data revealed that CCR7 disrupts late CD8^+^ T cell-DC interactions by “switching” the intracellular localization of the F-actin-promoting factor DOCK2 away from the IS.

**Figure 3.**
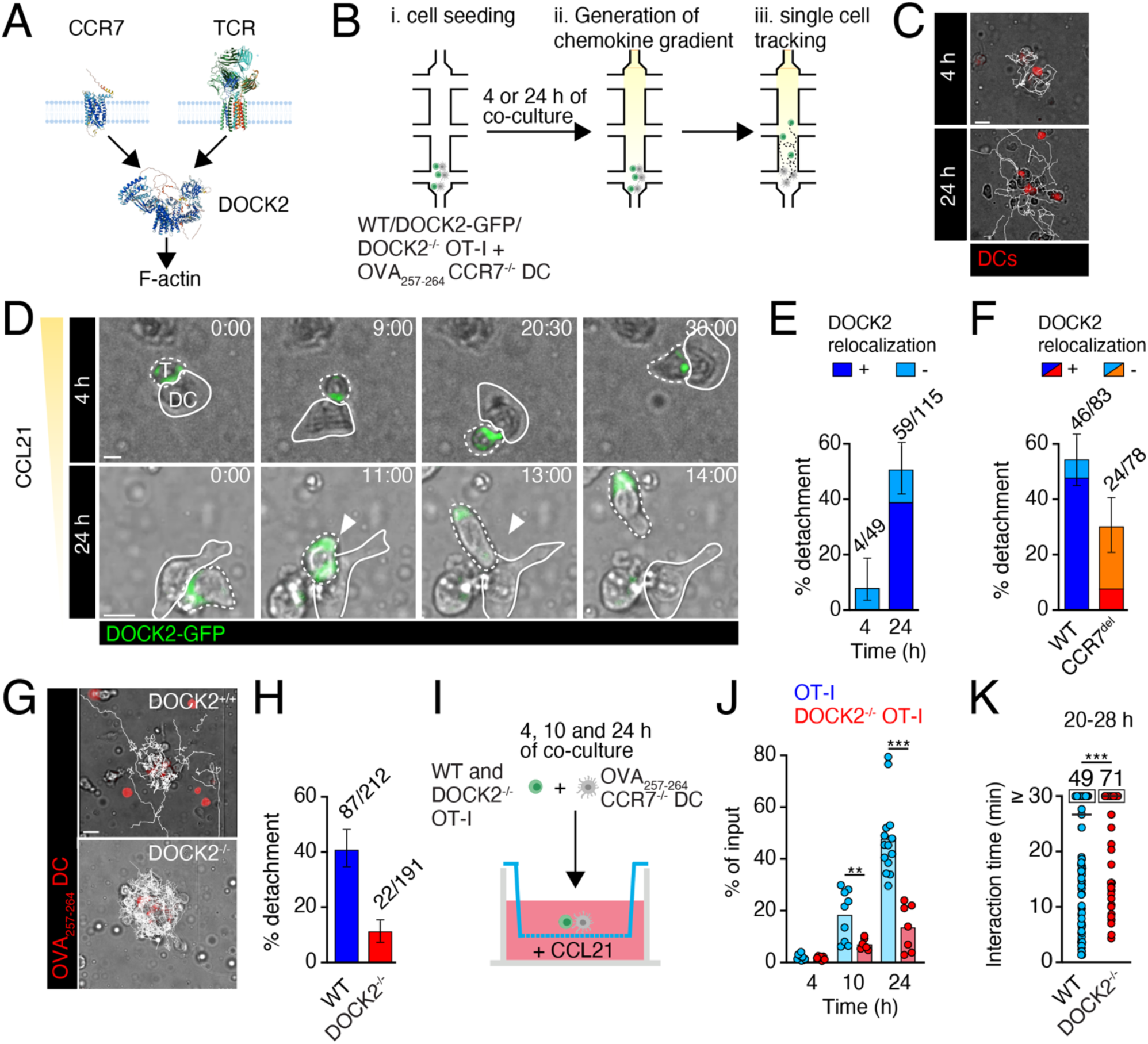
CCR7 signals reposition the Rac activator DOCK2 to promote CD8^+^ T cell detachment. **A.** Scheme of DOCK2 activation downstream CCR7 and TCR. Structures taken from AlphaFold (https://alphafold.ebi.ac.uk/). **B.** Layout of microfluidic chamber experiments. **C.** Example of OT-I clusters with OVA_257-264_-pulsed DCs after exposure to CCL21 at 4 or 24 h of interaction. White lines depict T cell tracks. Scale bar, 10 µm. **D.** Time-lapse images of DOCK2-GFP localization with respect to OVA_257-264_-pulsed DCs at 4 or 24 h of interaction. The arrowhead depicts T cell release with DOCK2 relocalization. Scale bar, 5 µm; time in min:s. **E.** Percentage of detachment events over total DOCK2-GFP OT-I-DC interactions with or without DOCK2 relocalization away from the IS. Numbers above bars indicate total detachment events over interacting cells. Error bars depict 95% confidence intervals for total detachment events. **F.** Percentage of detachment events over total CCR7^del^ DOCK2-GFP OT-I-DC interactions with or without DOCK2 relocalization away from the IS at 24 h. Numbers above bars indicate total detachment events over interacting cells. Error bars depict 95% confidence intervals for total detachment events. **G.** Time-lapse images of control and DOCK2^-/-^ OT-I T cells interacting with OVA_257-264_-pulsed DCs at 24 h of interaction. White lines depict T cell tracks. Scale bar, 15 µm. **H.** Percentage of detachment events over total WT and DOCK2^-/-^ OT-I T cell-DC interactions at 24 h. Numbers above bars indicate total detachment events over interacting cells. Error bars depict 95% confidence intervals for detachment events. **I.** Experimental layout for Transwell assays with DOCK2^-/-^ T cells. **J.** Chemotaxis of WT and 4/10/24 h OVA_257-264_-pulsed DC-interacting control or DOCK2^-/-^ OT-I T cells towards CCL21. **K.** Interaction times of control and DOCK2^-/-^ OT-I T cells with OVA_257-264_-pulsed DCs *in vivo*. Data in E, F, H, J and K are pooled from at least two independent experiments and analyzed using a Student’s *t*-test (J) or Mann-Whitney test (K). **, p < 0.01; ***, p < 0.001.

### Chemokine disruption of T cell-DC interactions safeguards acute and recall T_EFF_ responses

To examine the physiological impact caused by disruption of interacting T cell-DC pairs by lymphoid tissue chemokines, we separated OT-I T cells from OVA_257-264_-pulsed CCR7^-/-^ DCs after 10 or 24 h by exposing them to a CCL21 gradient. We then processed the resulting chemokine-detached (CD) OT-I T_EFF_ cohorts (CD-10 and CD-24, respectively) for bulk RNA sequencing (RNAseq). A third cohort was allowed to interact with peptide-pulsed DCs for 48 h, which is the minimal time commonly used in *in vitro* activation protocols for expansion of T cells for adoptive cell transfers (*59*, *60*), before separating them from DCs and processed for RNAseq (“non-detached”, ND) (**Figure 4A**). All three cohorts showed robust activation as assessed by similar numbers of differentially expressed genes (DEGs), including a downregulation of T_N_-expressed genes (e.g., *Klf2*, *S1pr1*) and an increase in effector gene transcription (e.g., *Il2ra*; **Figure S4A**). In principal component analysis (PCA) of transcriptomes of the timed cohorts, T_N_ clustered separately from CD-10 and CD-24 T_EFF_, which grouped closely together (**Figure 4B**). We also performed a bulk RNAseq analysis of non-migrated OT-I T cells (isolated from the top compartment of Transwell chambers) at the 24 h time point but did not observe a distinct clustering compared to migrated cells collected from the bottom chamber (**Figure 4B**). Notably, ND T_EFF_ clustered separately from both T_N_ and CD-10 and CD-24 T_EFF_ (**Figure 4B**). The segregation of the ND T_EFF_ versus the CD-10 or CD-24 transcriptome was further corroborated when comparing the regulation of the top 50 DEGs in the three cohorts (**Figure S4B**), and when performing a PCA of RNAseq samples isolated from all three cohorts at 60 h post stimulation (**Figure S4C**). Given the link between chronic TCR signaling and dysfunction, we investigated the expression of genes associated with impaired CD8^+^ T cell function between the three cohorts. While we did not detect distinct *Pdcd1* or *Ctla4* mRNA levels at these early time points, mRNA levels of *Nr4a* transcription factors (TFs), *Cd200*, *Lag3* and *Dapl1*, negative regulators of T cell activation, were significantly increased in ND but not CD-10 and CD-24 T_EFF_ (**Figure 4C**) (*61–63*). In addition, the TF *Bhlhe40*, which has been described to drive T cell exhaustion at the expense of stem-like T cell differentiation (*18*), was significantly increased in ND versus CD-24 T_EFF_. In contrast, effector function-related transcripts (*GzmA/B, Ccr2*, *Ccr5*) were decreased in ND versus CD-10 or CD-24 cohorts (**Figure 4C**). To correlate activation-induced gene expression profiles with protein levels in effector populations, we further cultured CD-10 and CD-24 T_EFF_ in the presence of unpulsed, activated DCs until 48 h. Afterwards, these cohorts, together with ND T_EFF_, were kept until 60 h post stimulation in the absence of DCs but with IL-2 supplementation to mimic continuous CD8^+^ T cell dwelling in lymphoid tissue. A proteome analysis (n = 6578 proteins analyzed) at 60 h post stimulation showed an increase in levels of the TFs Nr4a3, BACH2 and Rel in the ND cohort, whereas NFATc2, T-bet and Eomes were enriched in the CD-10 and CD-24 cohorts (**Figure 4D and Figure S5A**). These data suggested a delayed transition to an effector phenotype in the ND cohort, with a parallel increase in Nr4a TF levels associated with dysfunctional T cell activation and T cell exhaustion. Accordingly, we observed significantly increased levels of levels of Dapl1, PD-1, CD200 and LAG3 in ND as compared to CD-10 or CD-24 T_EFF_, whereas levels of granzyme (Gzm) B and C scaled inversely with DC interaction time (**Figure 4D and Figure S5A**). Using flow cytometry at 60 h post activation, we confirmed the inverse scaling of GzmB levels observed by proteome analysis, with highest abundance in the CD-10 T_EFF_ cohort. Furthermore, levels of PD-1, CD200 and LAG-3 were markedly increased in ND as compared to CD-10 and CD-24 T_EFF_, whereas CD200r and CTLA-4 levels were similar between all three cohorts (**Figure 4E**). A comparable GzmB and PD-1 expression pattern emerged when examining CD-10, CD-24 and ND CD8^+^ P14 T_EFF_ at 60 h post stimulation with LCMV gp_33-41_-pulsed DCs (**Figure S5B**). Finally, we examined whether mechanical separation of CD8^+^ T cells from TCR activation signals sufficed to reproduce immune checkpoint expression patterns. To this end, we used polyclonal stimulation with surface-bound anti-CD3ε and anti-CD28 mAbs (**Figure S5C**). While a 10 h stimulation was insufficient for OT-I T cell activation in this setting (not shown), extending the polyclonal activation period to 48 h yielded T_EFF_ with substantially increased levels of PD-1, CD200 and LAG3, and to a lesser extent, CD200r and CTLA-4 as compared 24 h-stimulated cells (**Figure S5D**). In sum, our findings suggest that chemokine-restricted T cell activation promoted a CD8^+^ T_EFF_ transcriptional program characterized by expression of cytotoxic factors such as GzmB. Extending stimulation to 48 h or beyond, the minimal time window commonly used in experimental and clinical T cell activation protocols (*60*), prevented or delayed expression of key cytotoxic factors, and led to a marked increase in factors associated with impaired effector functionality.

**Figure 4.**
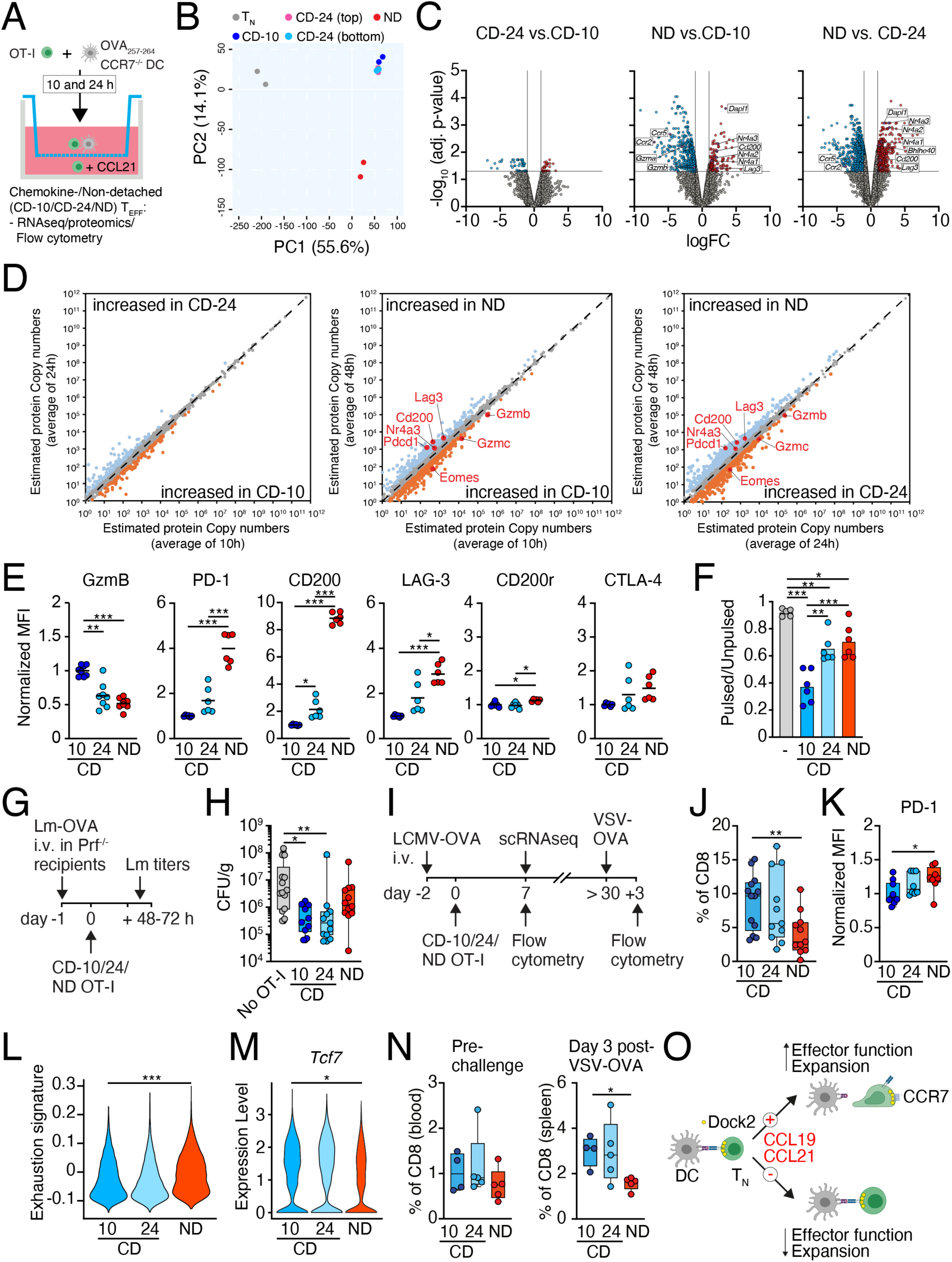
Stromal chemokine-triggered disruption of T cell-DC interactions safeguards CD8^+^ T_EFF_ differentiation. **A.** Experimental layout to generate 10 and 24 h “chemokine-detached” (CD-10 and -24, respectively) and “non-detached” (ND) OT-I T_EFF_. **B.** PCA of naive, CD-10, CD-24 and ND OT-I T cells immediately after separation from OVA_257-264_-pulsed DCs. Numbers in brackets indicate % of variance explained. **C.** Volcano plots of CD-10/CD-24/ND OT-I T_EFF_ cohorts with cutoff of logFC ≥ 1 and adjusted p-value ≥ 0.05. Selected genes are indicated. **D.** Protein copy numbers in CD-10, CD-24 and ND OT-I T cells at 60 h post stimulation. The dashed line represents proteins with the same copy numbers of the two indicated populations. Selected two-fold regulated proteins are highlighted with red. Average values of n = 4 biological replicates for CD-10 and CD-24, and of n = 3 biological replicates for ND T_EFF_ are shown. **E.** Flow cytometry analysis of CD-10, CD-24 and ND OT-I T_EFF_ at 60 h post stimulation. **F.** *In vitro* cytotoxicity of CD-10, CD-24 and ND OT-I T_EFF_. **G.** Experimental layout of Lm-OVA infection. **H.** Lm cfu in spleens of Prf^-/-^ recipients after adoptive transfer of CD-10, CD-24 and ND OT-I T_EFF_. **I.** Experimental layout of LCMV-OVA infection and recall response. **J.** Percentage of CD-10, CD-24 and ND OT-I T_EFF_ in spleens on day 7 p.i. **K.** Flow cytometry analysis of PD-1 expression on CD-10, CD-24 and ND OT-I T_EFF_ in spleens on day 7 p.i. **L.** Violin plots showing the exhaustion signature expression in CD-10, CD-24 and ND OT-I T_EFF_ on day 7 p.i. **M.** *Tcf7* expression in CD-10, CD-24 and ND OT-I T_EFF_ on day 7 p.i. **N.** Percentage of CD-10, CD-24 and ND OT-I T cells before (blood) and on day 3 post VSV-OVA rechallenge (spleen). **O.** Graphical summary of proposed model. Data are pooled from at least two independent experiments and analyzed using ANOVA with Dunnett’s post-test (E, F, H, J, K, L, M, N). *, p < 0.05; **, p < 0.01; ***, p < 0.001.

We next examined how lymphoid chemokine-triggered disruption of T cell-DC interactions affected T_EFF_ function. When exposed to peptide-pulsed target cells (**Figure S5E**), ND OT-I T_EFF_ displayed impaired *in vitro* cytotoxic activity compared to CD-10 T_EFF_ at 60 h post activation (**Figure 4F**). To assess the capacity to protect against microbial infection *in vivo*, we transferred CD-10, CD-24 and ND OT-I T_EFF_ at 60 h post activation into Lm-OVA-infected perforin-deficient (Prf^-/-^) mice, which have an impaired endogenous cytotoxic response (**Figure 4G**). Whereas both CD-10 and CD-24 OT-I T_EFF_ cohorts significantly lowered Lm levels by more than 10-fold as compared to “no OT-I” control animals, ND T_EFF_ failed to confer such protection (**Figure 4H**). A similar trend was observed when transferring T_EFF_ cohorts into WT recipients (not shown). This finding suggested that despite an initial delay in GzmB expression in the CD-24 T_EFF_ cohort, these cells became equally competent to control microbial infections as CD-10 T_EFF_. In contrast, ND T_EFF_ failed to exert potent antimicrobial activity at the early time points tested here.

Previous studies using varying pMHC-TCR affinities have uncovered an inverse correlation between the duration of *in vivo* T cell-DC interactions, with shorter interactions favoring accelerated effector function acquisition and T_EFF_ release from SLOs (*23*). Yet, despite an initial delay in acquiring cytotoxicity and egress from SLOs, high affinity pMHC-primed T_EFF_ eventually come to dominate the CD8^+^ T_EFF_ population during the late effector and memory phase of an immune response (*26*). To examine the phenotype of ND T_EFF_ at later time points of acute infection, we adoptively transferred CD8^+^ T cells into virus-infected hosts and examined expansion and phenotype on day 7 of the effector response (**Figure 4I**). At this time point, CD-10, CD-24 and ND T_EFF_ showed comparable frequencies of CD127^-^ KLRG-1^+^ short-lived effector cells (SLEC) versus CD127^+^ KLRG-1^-^ memory precursor effector cells (MPEC; **Figures S5F and S5G**). In contrast, significantly fewer ND OT-I T_EFF_ were recovered from day 7 p.i. spleens as compared to CD-10 and CD-24 T_EFF_ (**Figure 4J**), akin to a reduced recovery of T_EFF_ that had been stimulated with anti-CD3ε and -CD28 mAbs for 48 versus 24 h (**Figure S5H**). The recovered ND T_EFF_ maintained elevated PD-1 levels (**Figure 4K**). To characterize the three T_EFF_ cohorts in more detail, we performed a scRNAseq analysis of transferred cells isolated from day 7 LCMV-OVA-infected spleens. We identified 7 distinct clusters, ranked 0-6 by descending cell counts (**Figure S6A**). Cluster 0 contained cells enriched in stem-like markers (*Sell, Tcf7, Slamf6, Cd127, Id3, Myb*), while clusters 1, 2 and 4 contained cells expressing effector-like markers (*GzmA, GzmB, Cx3cr1, Itga4*). Cluster 5 showed an enriched type I IFN signature (*Ift1, Ifit3, Mx1*), whereas clusters 3 and 6 had elevated levels of cell cycling genes (*Hist1, Kif11, Mki67*). A further refinement of the functional signatures of the identified clusters uncovered that clusters 3 and 6 were enriched in an expression profile characteristic of exhausted T cells (**Figures S6C and S6D**) (*64*). In contrast, clusters 0 and 5 had a low signature associated with exhausted T cells (**Figure S6C**), reflecting the expression of markers of stem-like properties in CD8^+^ T cells (*65–68*). All three T_EFF_ cohorts were represented in all 7 clusters but in varying frequencies. Notably, ND T_EFF_ were more frequent in clusters 3 and 6, and least frequent in clusters 0 and 5 (**Figure S6B**). In accordance, ND T_EFF_ showed a significantly increased exhaustion signature (**Figure 4L**) and decreased *Tcf7* expression (**Figure 4M**) compared to CD-10 T_EFF_. Despite their lower numbers, ND T_EFF_ displayed an increased effector and cycling signature (**Figure S6E**). Finally, to address long-term preservation of T cell functionality, we examined the impact of T cell-DC interaction duration on recall response in the memory phase (**Figure 4I**). At > day 30 p.i., we observed comparable numbers of all three cohorts in blood (**Figure 4N**), suggesting that prolonged cell cycling partially compensates for decreased ND T_EFF_ numbers during acute infection. Nonetheless, we recovered fewer ND OT-I T cells from spleens after rechallenge with OVA-expressing vesicular stomatitis virus (VSV-OVA) as compared to CD-10 and CD-24 memory T cells (**Figure 4N**).

## Discussion

In sum, our data suggest that lymphoid tissue-secreted chemokines act as stromal immune checkpoints to restrict CD8^+^ T cell-DC interactions, revealing their dual function to enable clonal selection as well as to limit activation signal integration of responding CD8^+^ T cells (**Figure 4O**). Both processes are required to expedite the generation of cytotoxic T_EFF_ for rapid interception of microbial infections. Regulating responsiveness to stromal chemokines in interacting T cells allows activatory signal integration and transcriptional regulation to be adjusted in a fast, reversible, and individual manner, and bypasses the need for regulation by nuclear events, which take much longer to manifest (e.g., through *de novo* expression of immune checkpoint inhibitors), or by signals that might influence adjacent cells (e.g., diffusible cytokines). The observation that chemokine-driven disruption of DC engagement expedites the generation of cytotoxic T_EFF_ and safeguards their functionality throughout the entire immune response underscores the importance of factors produced by the tissue microenvironment in which T_N_ activation has evolved. Our data further identify a critical time window between 24 and 48 h of continued T cell-DC interactions, where a marked increase in expression of immune checkpoints associated with CD8^+^ T cell dysfunction, such as PD-1, becomes evident (*69–72*). Besides immune checkpoints, prolonged T cell-DC interactions conceivably affect multiple aspects of T_EFF_ function, including epigenetic imprinting, metabolic activity, and pathways regulating cell cycling and survival. While the precise mechanism underlying ND T_EFF_ dysfunction remains unclear, it was striking to note the increased levels of Nr4a TFs in this population compared to CD-10 and CD-24 T_EFF_. Expression of Nr4a members has been linked to a decreased accumulation of stem-like TCF1^+^ cells and the development of T cell dysfunction at late time points of an immune response (*61*, *63*, *73*), which might contribute to the lower recovery and decreased frequency in stem-like markers of adoptively transferred ND T_EFF_ observed here. Dysfunctional T cell states have been observed early after tumorigenesis and viral infections (*20*, *70*, *71*, *74–76*). In this regard, our observation of prolonged T cell priming in the absence of a CCR7 “disruptor” points to a scenario where low lymphoid tissue chemokine levels in chronically inflamed or tumor-draining LNs (*40–42*) contribute to transcriptional imprinting of a dysfunctional T cell differentiation state, which becomes epigenetically fixed under persistent high Ag loads (*67*, *77*, *78*). Our data are also consistent with the observation that T_EFF_ generated early in chronic infection models, i.e., prior to the downregulation of CCR7 ligands in lymphoid tissue stromal cells, resemble T_EFF_ formed during acute infection (*75*). Physiological processes are subject to multiscale control systems, and each cell integrates hundreds of feedback loops to adapt its functionality (*79*). Accordingly, adaptive immune reactions are controlled by myriads of – often incompletely characterized – feedback systems that ensure tailored yet non-overshooting responses. Our findings uncover a stromal chemokine checkpoint-driven detachment constitutes a fundamental feedback system for optimal T cell activation. Paracortical stromal cells have been long recognized to shape adaptive immunity (*6*, *80*, *81*). Focusing on antimicrobial CD8^+^ T cell responses, we expand their functions to include the capping of T cell priming by chemokines before transcriptionally regulated immune checkpoints fulfill this function in effector and memory T cells (*30*, *82*). On a translational level, designing protocols with shorter, and therefore more physiological, T cell activation duration prior to adoptive transfer into human patients might contribute to increased functionality and long-term persistence of T_EFF_ for long-lasting cancer immunotherapy. In support of such an approach, the presence of CCL21 and ICAM-1 during *in vitro* CD8^+^ T cell priming equips T_EFF_ with increased antitumoral activity (*83*). In sum, our data identify an unexpectedly short, critical time window for optimal T cell activation, which requires input by lymphoid stromal chemokine signals before “classical” immune checkpoints regulate TCR signaling.

## Materials and Methods

### Mice

OT-I TCR transgenic mice (*22*) on a wild type or RAG1^-/-^ background were backcrossed to hCD2-dsRed (“DsRed”) (*84*), Nur77^GFP^ (*21*), Tg(UBC-GFP)30Scha (“GFP”) (*85*) and tdTomato-expressing Ai14 x ZP3 mouse lines (“tdTom”) (*86*, *87*). OT-I mice were further crossed with CCR7^-/-^ (*88*), DOCK2^-/-^ (*53*) or DOCK2-GFP (Dock2^tm1Ysfk^) mouse lines (*58*), respectively. C57BL/6 mice were purchased from Janvier (AD Horst). All mice were maintained at the animal facility of the University of Fribourg. All animal work has been approved by the Cantonal Committee for Animal Experimentation and conducted according to federal guidelines.

### Adoptive cell transfer and infections

CD8^+^ T cells were negatively isolated from spleen and peripheral LNs of donor mice using the Mojosort^TM^ Mouse CD8^+^ T cell Isolation Kit (BioLegend). T cell purity was confirmed to be > 95% by flow cytometry. DCs were generated from C57BL/6 bone marrow by culturing for 7 days in complete medium (RPMI 1640 (Gibco, 21875-034) containing 10% FCS (Gibco, 10270-106), 10 mM HEPES (P05-01100), 100 U/mL penicillin and 0.1 mg/mL streptomycin (P06-07100), 2 mM L-glutamine (P04-80100), 1 mM sodium pyruvate (P04-43100) and minimum essential medium essential amino acids (P08-32100; from PAN Biotech) supplemented with 10% culture supernatant of SP2/0 cells expressing Flt3-ligand, followed by activation with 0.5 µg/mL *S. typhimurium*-derived LPS (Sigma) for 24 h.

For IVM experiments, we i.v. transferred 3 x 10^6^ T cells/mouse (in co-injections 1.5 x 10^6^ of each population) 18 h post s.c. DC vaccination (1 x 10^6^ DCs in 10 µL/foot hock). For CCR7^-/-^ OT-I cells, 3-4 x 10^6^ CCR7^-/-^ and 1.5 x 10^6^ control OT-I cells were injected to obtain comparable numbers in LNs. We injected 100 µg anti-mouse L-selectin antibody i.p. (Mel-14, BioXcell, BE0021) 3 h post T cell transfer to prevent further homing and to synchronize the initiation of T_N_-DC interaction. For Lm-OVA clearance experiments, we transferred 1 x 10^5^ CD-10, CD24 or ND OT-I T cells per mouse. In some experiments, mice received 2 x 10^6^ pfu LCMV clone 13 i.p. 3 d prior to DC transfer. For microbial infections, mice received 2 x 10^6^ Nur77^GFP^ OT-I or 1 x 10^5^ CD-10, CD24 or ND OT-I T cells followed by 10^5^ p.f.u. LCMV-OVA i.p. (*89*) or 5 x 10^3^ cfu *Listeria monocytogenes*-OVA (Lm-OVA) i.v. as described (*90*). For rechallenge experiments, we administered 2 x 10^6^ VSV-OVA i.p. in LCMV-OVA memory recipients.

### Chemotaxis

Activated DCs were pulsed with 100 nM peptides (EMC microcollections, Tübingen, Germany) for 45 min. For intravital imaging, DCs were stained with 20 µM CellTracker blue^TM^ (ThermoFisher) for 20 min and injected s.c. into the foot hock of recipient mice (1 x 10^6^ DCs). Isolated OT-I were co-cultured with DCs from CCR7^-/-^ mice in Transwell chambers (5 µm pore size) and exposed to 50 nM CCL21 (Peprotech, 250-13), CCL19 (R&D, 440-M3), CXCL10 (R&D, 466-CR) or CCL3 (Peprotech, 250-09) for 1 h before collecting cells from the lower well. For some experiments, migrated OT-I cells were cultured with non-peptide pulsed DC from the same culture until 48 h. Isolation of OT-I T cells and DCs at 48 h of co-cultures was performed by incubation with 10 µg/mL anti MHC-I (H-2) antibody (BioXcell, BE0077) for 1 h before performing negative CD8 isolation. For mAb-triggered T cell activation, 0.3 x 10^6^ OT-I T cells were placed on flat-bottom 96 well plates coated with 5 µg/mL anti-CD3ε (clone 145-2C11; Biolegend) and stimulated with 1 µg/mL anti-CD28 mAb (clone 37.51, BioLegend) in solution. To stop TCR stimulation, OT-I cells were carefully transferred into uncoated wells. Plates were microscopically examined to confirm the complete transfer of OT-I cells to the adjacent well.

### Under agarose assays

A circle of 17-mm diameter was cut into the center of 60-mm dishes and the hole was sealed using a 24-mm glass coverslip and aquarium silicone (Marina). After drying, we overlaid a 5 mm high ring cut from a 15 mL tube and sealed it with paraffin. Coverslips were coated with 20 µg/mL Protein A (BioVision, 6500-10) for 1 h at 37°C, washed three times with PBS and blocked with 1.5% BSA for 1 h at 37°C. Cover glasses were coated with 100 nM recombinant ICAM-1 Fc (R&D, 796IC) for 2 h at 37°C before being washed twice with PBS. Coverslips were blocked with 1.5% BSA for 1 h at 37°C before addition of 500 nM CCL21 (Peprotech, 250-13) in cell culture medium. For agarose overlay, 10 mL 2X RPMI containing 20% FCS and 5 mL of 2X HBSS were heated up to 56°C in the water bath. 100 mg SeaKem Gold agarose (Lonza, 50152) was heated in 5 mL MilliQ water and added to the prewarmed medium to obtain a 0.5% agarose mix. After cooling down to 37°C, the CCL21 solution was removed and 500 µL of the agarose was added on the coverslip. The agar was left to solidify for 5 min at RT and subsequently 20 min at 4°C. T cell-DC conjugates were co-cultured until 4 h or 24 h and gently injected under the agarose in a volume of approximately 2-5 µL directly before imaging. Timelapse images were taken with a DeltaVision Elite widefield fluorescent microscope equipped with a 20X, NA 0.75 objective (Olympus). Images were taken from 8-10 FOV every 20 s for 60 min. Data were analyzed using Imaris (Bitplane).

### Flow cytometry

LNs and spleens were harvested at indicated timepoints and single cell suspensions were generated using cell strainers (70 µm mesh size). Fc receptors were blocked using 2.5 µg/mL anti-CD16/32 (Biolegend, 101330) for 20 min in PBS containing 5 mM EDTA and viability dyes Zombie Green or Red (R&D). Following antibodies for flow cytometry were used: CD8a-APC-Fire750 (Biolegend, 100766), CD45.1-PE (Biolegend, 110708), CD279/PD-1-PE clone (Biolegend, 29F.1A12), CCR7-biotin (eBiosciences, 13-1971-85), CCR7-PE (Biolegend, 120105), CXCR3-BV605 (Biolegend, 126523), CCR5-APC (Biolegend, 107019), CD223/Lag3-BV711 (BioLegend 125243); CD200/OX2-PE/Cy7 (BioLegend 123818); CD152/CTLA4-PE (BioLegend 106305); CD200R/OX2R-FITC (BioLegend 123909), Propidium Iodide (ThermoFisher) and CD69-PE (Biolegend, 104508), followed by fixation with 4% paraformaldehyde (Electron Microscopy Sciences) for 20 min on ice. For intracellular labeling of GzmB, we used Cytofix/Cytoperm^TM^ kit (20 min fixation on ice; BD), followed by GzmB-AF647 (Biolegend, 515405) and washing 1x with Perm/Wash^TM^ buffer (BD).

### Intravital microscopy

C57BL/6 mice were anesthetized with ketamine/xylazine/acepromazine and the right popliteal LN was surgically exposed without damaging the organ capsule or lymph vessels as described (*23*). Prior to imaging, AF633-conjugated MECA-79 (10 μg/mouse) was injected i.v. to visualize high endothelial venules. Intravital imaging was performed with an Olympus BX50WI fluorescence microscope equipped with a 25X objective (Nikon, NA 1.1) and a TrimScope 2PM system controlled by ImSpector software (LaVision Biotec). Timelapse image series were acquired using an automated tissue-drift correcting software (*91*). A Ti:sapphire laser (Mai Tai HP) was tuned to 780 or 840 nm. Image stacks were transformed into volume-rendered four-dimensional videos with Imaris (Bitplane), which was also used for semi-automated tracking of cell motility. Interaction times were determined by manual inspection.

### Microfluidic chamber experiments

Chip design, fabrication and set-up were described before (*92*). WT, DOCK2^-/-^ or DOCK2-GFP OT-I T cells were mixed with activated OVA_257-264_-pulsed DC in a1:2 ratio of T cells to DCs and loaded into the microchamber. After 4 or 24 h, a chemokine gradient was established through diffusion by alternatingly opening valves on the top and bottom of the chamber which were connected to media lines with either vehicle (sink) or 500 nM CCL21 (chemokine source; Peprotech, 250-13). Whole chambers were imaged for a duration of 2-4 h using a Nikon Ti2E equipped with a Photonics Prime 95B camber and a 40X Plan Apo Lambda Objective, NA 0.95.

### CRISPR/Cas-deletion of CCR7

CCR7 crisprRNAs (crRNA) were designed using DESKGEN online tool (www.deskgen.com). Alt-R^®^ CRISPR-Cas9 crRNA (custom design), trans-activator RNA (tracrRNA) (1072534) and Alt-R^®^ CRISPR-Cas9 negative crRNA (226567203) were purchased from Integrated DNA Technologies (Coralville, IA) and reconstituted at 100 µM with nuclease free duplex buffer (Integrated DNA Technologies). One µL each of crRNA and tracrRNA were annealed to form guide (g) RNA at 95°C for 5 min using a thermal cycler and cooled to room temperature. Annealed gRNA was mixed with TrueCut Cas9 v2 (A36499, Thermo Fisher) at a ratio of gRNA: Cas9 = 1.8 µL: 1.2 µL (equivalent to 90 pmol : 36 pmol) and left at room temperature for > 10 min to generate ribonuclein protein (RNP) complexes. Naive DOCK2-GFP OT-I T cells were resuspended in P4 Primary Cell 4D-Nucleofector™ X Kit S (V4XP-4032, Lonza) buffer solution at a cell concentration of 5 x 10^6^ cells in 20 µL, mixed with RNP complex solution and added to Nucleocuvette™ strip well. Three RNP complexes in 9 µL were used per reaction. Cells were nucleofected using a 4D-Nucleofector™ with X-Unit (Lonza) and pulse DS137. After nucleofection, 100 µL of pre-warmed complete medium containing 20 ng/mL IL-7 (402-ML-020/CF, R&D Systems) was added to each Nucleocuvette™. OT-I T cells were gently mixed by pipetting and aliquoted into a flat-bottom 96-well plate and cultured in a total volume of 200 µL complete medium containing IL-7 at 2 x 10^6^ cells for 8 days at 37°C, 5% CO_2_. On day 4 post nucleofection the medium was partially replaced. The three crRNA sequences used in this study are as follows: 5’-CATCGGCGAGAATACCACGG-3’, 5’-ACGCAACTTTGAGCGGAACA-3’, 5’-CCTGGACGATGGCTACGTAG-3’. Nucleofected OT-I T cells were cultured in RPMI1640 medium supplemented with 10% FCS, 2 mM L-glutamine, 0.1 mM non-essential amino acids, 1 mM sodium pyruvate, 100 U/mL penicillin, 0.1 mg/mL streptomycin, 50 µM 2-mercaptoethanol, 10 mM HEPES in the presence of 20 ng/mL IL-7 (R&D, 402-ML-020/CF,) at 37°C, 5% CO_2_. On day 8, cells were harvested and stained with PE-conjugated anti-mouse anti-CD197 (CCR7) (Biolegend, clone 4B12). After two rounds of wash, cells were resuspended in staining buffer containing 0.5 µg/mL propidium iodide. OT-I cells nucleofected with CCR7-RNP were sorted for viable CCR7^del^ fraction, and control cells we sorted for viable fraction using a FACSAria™ Fusion cell sorter (BD Biosciences). Sorted cells were collected in complete medium containing 5 ng/ml IL-7.

### Bulk RNA sequencing

Migrated or beads-separated RAG^-/-^ OT-I T cells were processed using the ReliaPrep^TM^ RNA Cell Miniprep System (Promega #Z6011) and total RNA was isolated following the manufacturer’s instructions. DNA digestion was performed with TURBO DNA-free Kit (Invitrogen #AM1907). RNA was quantified with Qubit^TM^ RNA HS Assay Kit (Invitrogen # Q32852) and RNA integrity was assessed with the Agilent RNA 6000 Pico Kit (Agilent #5067-1513) on a Bioanalyzer instrument (Agilent). RNA samples were processed with the “SMART-seq Ultra Low Input 48” library protocol in order to obtain 33.0M clusters of fragments of 1 x 75 nt of length through NextSeq 500 High 75. Raw reads were aligned to mouse genome build GRCm38 using STAR aligner (*93*). Gene counts were generated using featureCounts (part of the Subread package,(*94*)) based on GENCODE gene annotation version M22. To discard genes highly expressed by one sample only, a filter of cpm >= 2 in at least two samples was added. Read counts were normalized with the Trimmed Mean of M-values (TMM) method (*95*) using calcNormFactors function and then Voom (*96*) was applied.

### Data-independent mass spectrometry (DIA-MS) for proteome analysis

CCL21-migrated and beads-separated OT-I T_EFF_ at 60 h post activation were isolated as above and their pellets frozen. Cell pellets were lysed by SDC buffer (1% sodium deoxycholate in 50 mM ammonium bicarbonate buffer, pH 8.5), proteins were reduced with 2 mM dithiothreitol for 30 min at room temperature and alkylated with 10 mM iodoacetamide in dark for 30 min at RT, followed by trypsin digestion o.n. at 37 °C. The tryptic peptides were desalted by STAGE-tips and dried by speed Vac. The dried peptides were redissolved in buffer A (0.1% formic acid in water) and separated by EasyLC 1000 nanoflow-HPLC system (ThermoFisher Scientific) with a 20 cm fused silica column with an inner diameter of 75 μm and in-house packed with C18 (ReproSil-Pur 120 C18-AQ, 1.9 μm, Dr. Maisch). Peptides were loaded with solvent A at a max. pressure of 800 Bar and eluted with a step gradient of solvent B (0.1% formic acid in 80% acetonitrile) from 5% to 30% within 85 min, from 30% to 60% within 6 min, followed by increasing to 100% in 2 min at a flow rate of 250 nL/min. MS/MS analysis was performed on a nano-electrospray ion source equipped QExactive plus mass spectrometer (ThermoFisher Scientific). The spray voltage was set to 2.3 kV with a capillary temperature of 250°C. Mass spectrometer was operated in positive polarity mode and MS data were acquired in a data-independent mode (DIA-MS). The MS spectra (350-1’200 m/z) were acquired at a resolution of 140’000, an max. injection time (MIT) of 60 ms and automatic gain control (AGC) target value of 3 x 10^6^. The precursors were fragmented using a stepped normalized collisional energy (NCE) of 25.5, 27, and 30 by higher-energy collisional dissociation (HCD). MS/MS scans were acquired with a resolution of 35’000, AGC target value of 3 x 10^6^ and MIT was selected as automatic mode, isolation window of 31.4 m/z. MS raw files were analyzed using spectral library-free DirectDIA analysis approach by Spectronaut V17 (Biognosys AG) (*97*), data were searched against Uniprot full-length Mus musculus database (22’136 entries, released April, 2016). Cysteine carbamidomethylation was set as a fixed modification, protein amino-terminal acetylation and methionine oxidation were set as variable modification. Calibration was set to system default. Identification was performed using precursor PEP cutoff of 1 and precursor and protein Q-value cutoff of 1%. For quantification, interference correction was enabled with at least two MS1 and three MS2 spectra. Quantity was determined on MS2 level using areas of XIC peaks and cross run normalization was enabled. The Spectronaut results were analyzed using Perseus (Version 2.0.3.0) (*98*). Protein copy number estimations were calculated as described (*99*).

### Single cell RNA sequencing

CD-10, CD-24 and ND OT-I T cells were adoptively transferred into day 2.5 LCMV-OVA-infected mice and sorted from spleens at day 7 p.i. for single cell RNAseq. Single cell data analysis was performed using Seurat (v4.0.2) (*100*). Cells with sufficient bioinformatic quality were obtained after applying a filter of at least 200 genes expressed per cell and only genes expressed in at least 3 cells were retained. Moreover, cells with more than 10% of reads mapped to mitochondrial genes were also excluded from the analysis. We also removed one cluster with high expression of *Gm42418* and *AY036118* genes associated with ribosomal RNA contamination. UMI count matrix was further normalized and scaled following the standard Seurat workflow and UMAP dimensionality reduction was then applied on first 30 Principal Components after running PCA. Unbiased clustering was computed using the FindClusters function in Seurat with default parameters and a resolution value of 0.3. Specific markers for the different unbiased clusters were found using the function FindAllmarkers or FindMarkers in Seurat with default parameters. Violin plots of specific genes were produced with VlnPlot Seurat function. The exhaustion signature was created from the chronic “Tcf1-” signature by using the 100 most significantly increased genes in this population (*101*) and was calculated with the AddModuleScore function in Seurat. The effector and cycling signatures were from (*102*).

### In vitro killing assay

RAG^-/-^ OT-I T cells were chemokine-separated at 10 and 24 h from OVA_257-264_-pulsed DCs as above and cultured with non-pulsed DCs until 48 h. After magnetic bead separation, cells were cultured in complete medium supplemented with rmIL-2 (10 ng/mL; R&D, 402-ML) until 60 h. B cells were isolated from spleens of C57BL/6 mice (MojoSort Mouse Pan B cells, 480052) and labeled with CellTrace violet (5 µM; Thermofisher, C34557) for 20 min at 37°C. After washing with cell culture medium, B cells were pulsed with 500 nM OVA_257–264_ peptide for 45 min at 37°C, washed and mixed 1 : 1 with unpulsed B cells. 5 x 10^4^ OT-I T cells for each condition were mixed with 1 x 10^5^ mixed B cells for 15 h in 96 well plates. Viability was measured by Zombie Green (BioLegend) staining at the LSR Fortessa II (Beckton Dickinson).

### Statistical analysis

Two-tailed, unpaired Student’s *t*-test, Mann-Whitney *U*-test, one-way ANOVA with Dunn’s multiple comparisons test or Kruskal-Wallis test was used to determine statistical significance (Prism, GraphPad). Whiskers in “box and whisker” plots depict a range of 90-100% of individual values with values exceeding the range shown as individual dots) while the box comprises 50% of all data points and the line within the box indicates the median. Significance was set at *p* < 0.05.

## Acknowledgements

We thank Federica Conedera and Sergio Sanchez Luquin for help with Ca^2+^ flux analysis, Pamela Nicholson and Remy Bruggmann (University of Bern, Switzerland) for help with RNA sequencing, and Michael Sixt (ISTA, Austria) and Thorsten Mempel (Harvard University, Boston, MA) for critical reading of the manuscript. The microfluidic chips were kindly provided by Tim Schröder and Philip Dettinger (ETH, Zürich, Switzerland). This work benefitted from the BioImage Light Microscopy Facility and Cell Analytics Facility of the University of Fribourg.

## Author contributions

LMA, DCC, PD, JZ, JBdA, PP, JMM, JA: Investigation and methodology

CL, MK, CK, AHG: Bioinformatic analysis

MM, JD, MI, JVS: Supervision and funding aqcuisition

LMA, JVS: Conceptualization and writing

## Data availability

The mass spectrometry proteomics data have been deposited to the ProteomeXchange Consortium via the PRIDE partner repository (*103*) with the dataset identifier PXD049320: Username; Password: v7No1pAO). The RNAseq data for this study have been deposited in the European Nucleotide Archive (ENA) at EMBL-EBI under accession number PRJEB74611 (https://www.ebi.ac.uk/ena/browser/view/PRJEB74611).

## Supplemental Figures

**Figure S1.**
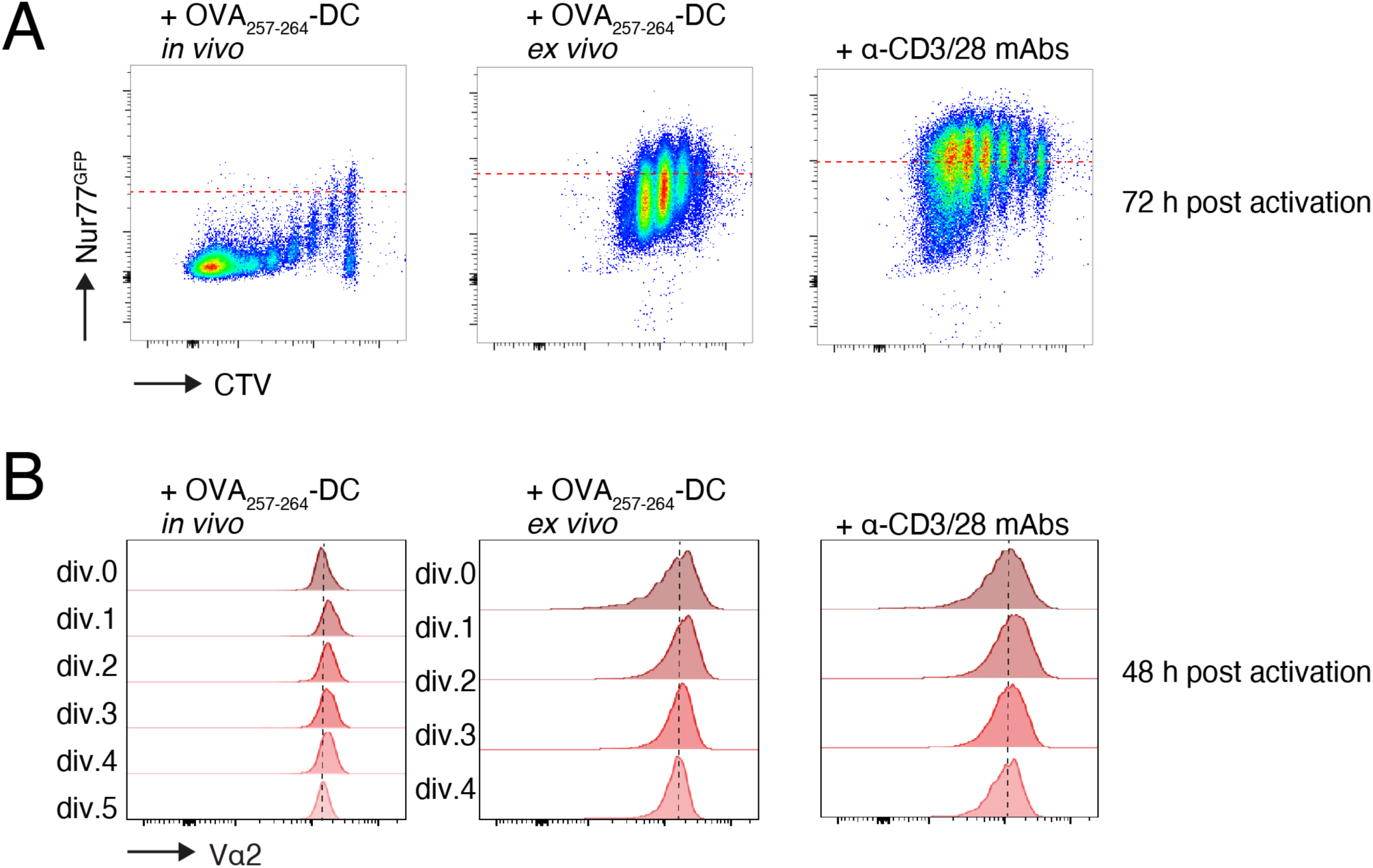
TCR signaling during *in vivo* and *in vitro* activation. **A.** Flow cytometry of CTV-labeled Nur77^GFP^ OT-I T cells at 72 h post activation. Red dotted lines depict GFP expression in activated, non-divided OT-I T cells. **B.** TCR surface levels during *in vivo* and *ex vivo* activation. Data are representative of at least two independent experiments.

**Figure S2.**
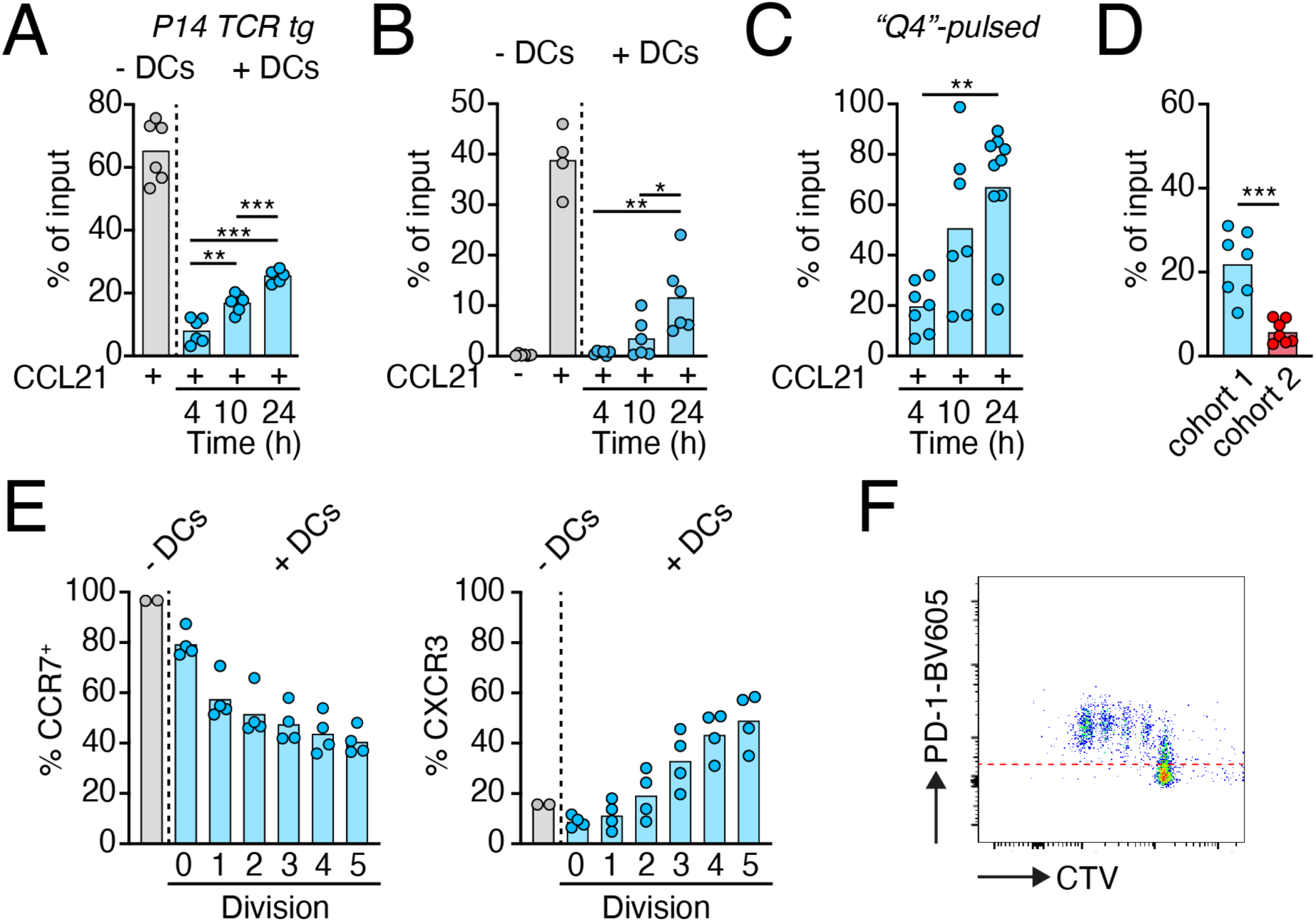
CCR7 ligands disrupt T cell-DC interactions. **A.** Chemotaxis of P14 T cells left unstimulated or interacting for 4, 10 or 24 h with LCMV gp33_33-41_-pulsed DCs towards CCL21. **B.** Chemotaxis of OT-I T cells left unstimulated or interacting for 4, 10 or 24 h with OVA_257-264_-pulsed CCR7^+/+^ DCs towards CCL21. **C.** Chemotaxis of OT-I T cells interacting for 4, 10 or 24 h with Q4-pulsed DCs towards CCL21. **D.** Chemotaxis of OT-I T cells interacting for 24 h (cohort 1) and 4 h (cohort 2) with OVA_257-264_-pulsed DCs towards CCL21 in same Transwell chamber. **E.** Flow cytometry analysis of CCR7 and CXCR3 expression on CTV-loaded OT-I T cells isolated from OVA_257-264_-pulsed DC-containing LNs at 48 h post T cell transfer. **F.** Flow cytometry of PD-1 expression on CTV-loaded OT-I T cells isolated from OVA_257-264_-pulsed DC-containing LNs at 48 h post T cell transfer. Red dotted line indicates FMO. Data in A-E are pooled or representative (F) from at least two independent experiments and analyzed using ANOVA with Dunnett’s post-test (A, B, C) and Student’s *t*-test (D). *, p < 0.05; **, p < 0.01; ***, p < 0.001.

**Figure S3.**
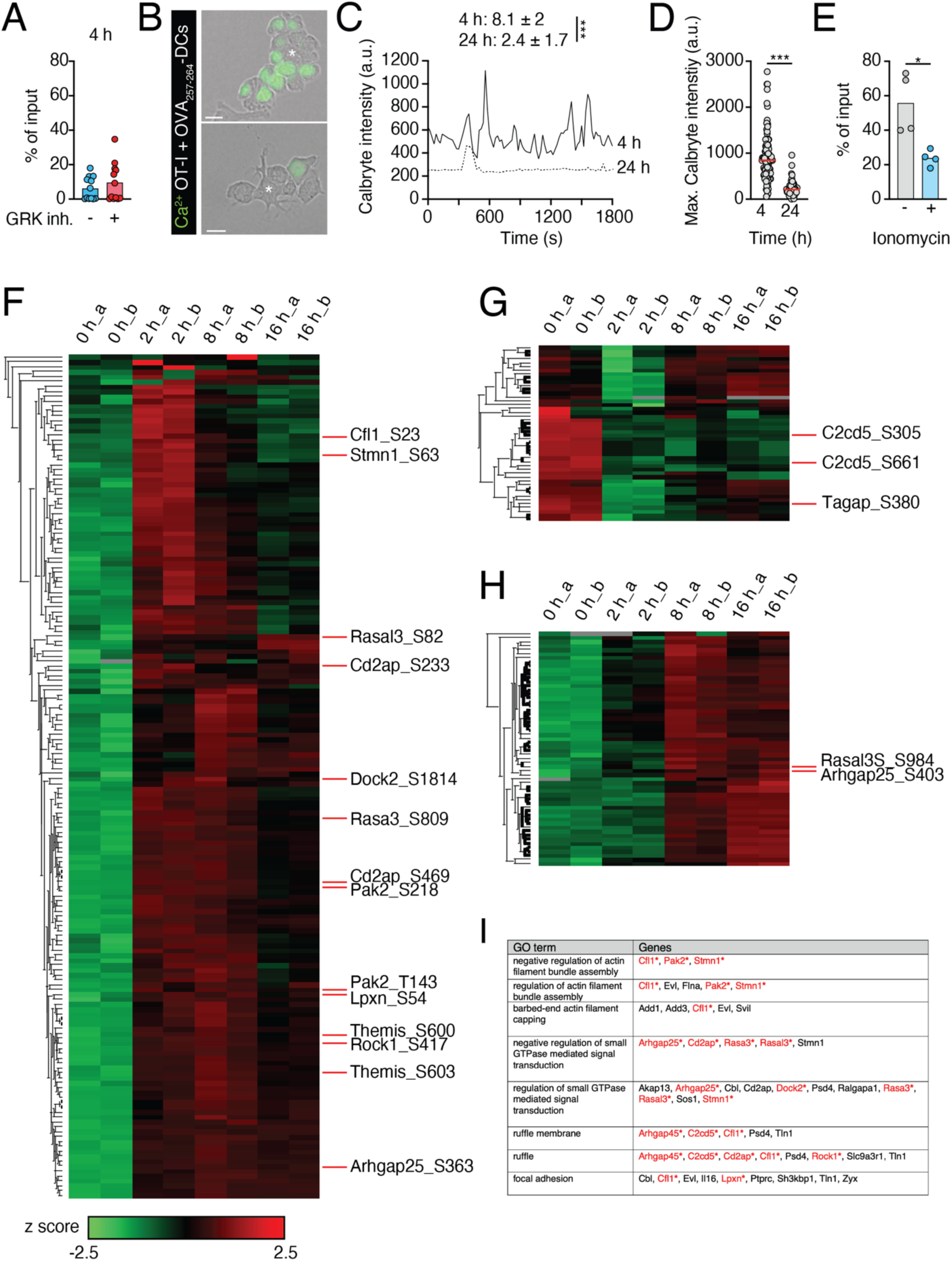
Analysis of signaling pathways governing CD8^+^ T cell responsiveness to chemokines. **A.** Chemotaxis of control and pan-GRK-inhibited OVA_257-264_-pulsed DC-interacting OT-I T cells towards CCL21 at 4 h of interactions. **B.** Image of OVA_257-264_-pulsed DC co-culture with Calbryte-loaded OT-I T cells at 4 or 24 h of interactions. Asterisks depict DCs. Scale bar, 5 µm. **C.** Calbryte intensity in selected OT-I T cells. Numbers depict median Ca^2+^ fluxes (± SD) over 30 min in n = 100 cells. **D.** Maximum Calbryte intensity at 4 and 24 h of interactions (n = 100 cells). **E.** Chemotaxis of OT-I T cells after 4 h of ionomycin treatment. **F.** Heatmap of early phosphorylated proteins after mAb-stimulated T cell activation. **G.** Heatmap of dephosphorylated proteins after mAb-stimulated T cell activation. **H.** Heatmap of late phosphorylated proteins after mAb-stimulated T cell activation. **I.** GO terms of TCR-triggered phosphorylated proteins involved in cytoskeletal regulation. Data in A and E are pooled from two independent experiments and in B-D from one experiment with 15 FOVs and analyzed using an unpaired Student’s *t*-test (A, C, D, E). Original data in F-I are from (*52*).

**Figure S4.**
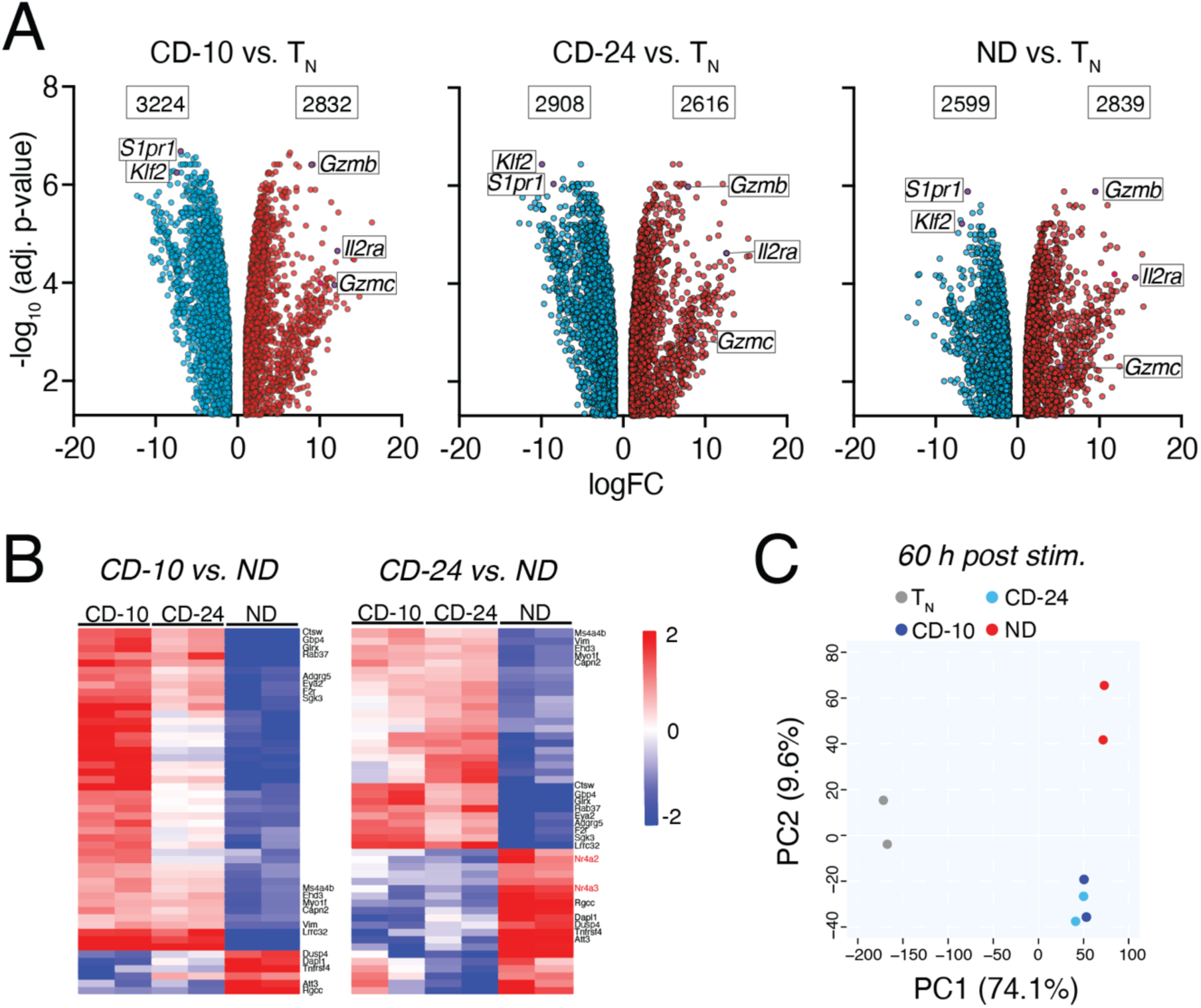
Transcriptome of chemokine-detached and non-detached T_EFF_. **A.** Volcano plots of DEGs in CD-10/CD-24/ND OT-I T_EFF_ versus T_N_ immediately after separation from OVA_257-264_-pulsed DCs. Only DEGs with cutoff of logFC ≥ 1 and adjusted p-value ≥ 0.05 are depicted, with numbers of up- or downregulated DEGs shown in box. Selected genes are indicated. **B.** Heat map of 50 most DGEs between CD-10 versus ND and CD-24 versus ND OT-I T_EFF_ immediately after separation from OVA_257-264_-pulsed DCs. Common genes between both comparisons are indicated, together with Nr4a genes in the CD-24 versus ND cohort. **C.** Principal component analysis of naive, CD-10, CD-24 and ND OT-I T cells at 60 h post stimulation. Numbers in brackets indicate % of variance explained.

**Figure S5.**
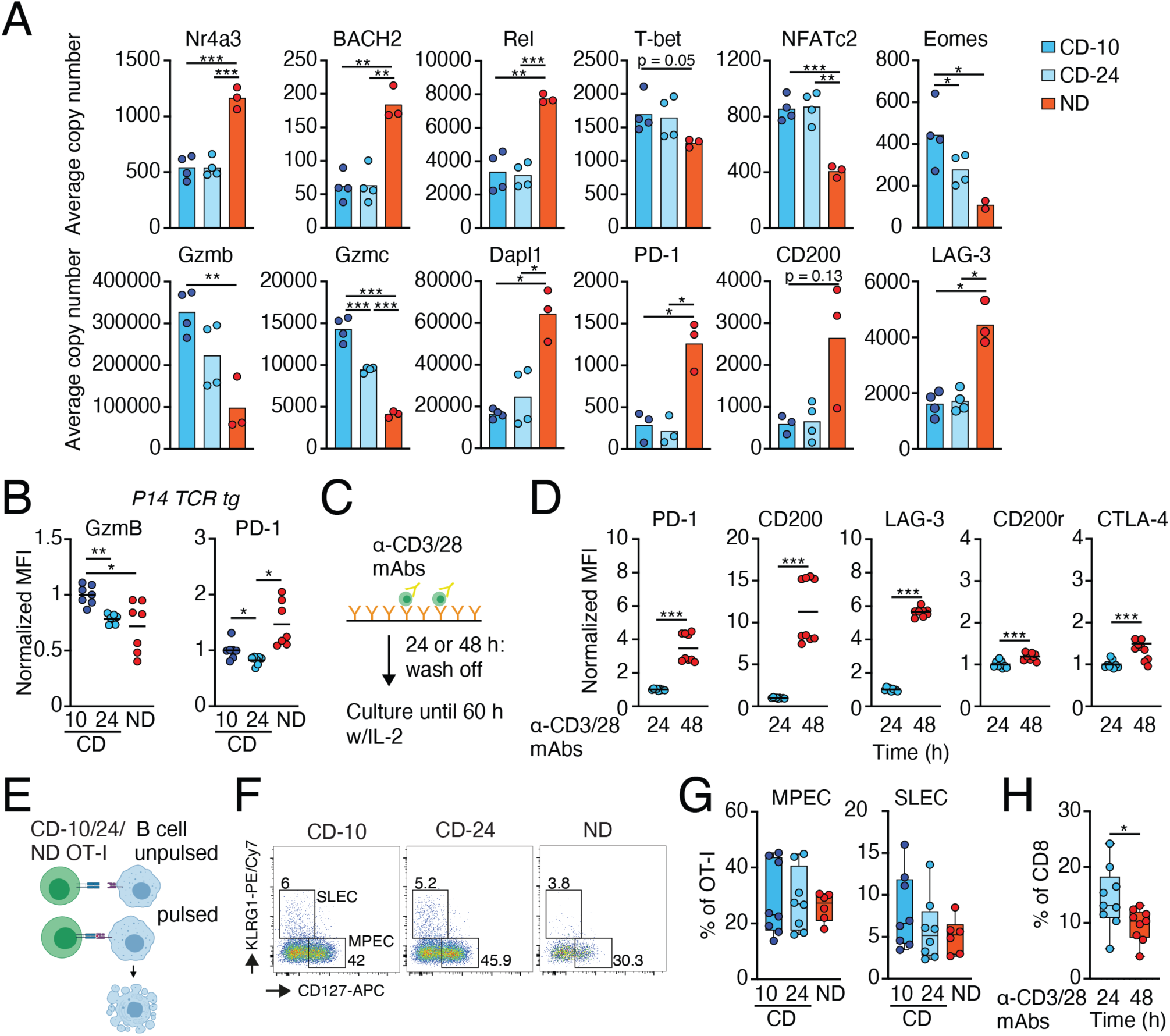
Effector differentiation of chemokine-detached and non-detached T_EFF_. **A.** Average copy numbers of selected proteins in CD-10, CD-24 and ND OT-I T_EFF_ at 60 h post stimulation. **B.** Flow cytometry analysis of CD-10, CD-24 and ND P14 T_EFF_ activated with LCMV gp_33-41_-pulsed DCs at 60 h post stimulation. **C.** Experimental layout for mAb-mediated T cell activation. **D.** Flow cytometry analysis of 24 and 48 h-mAb stimulated CD8^+^ T cells. **E.** Experimental layout of *in vitro* T_EFF_ function. **F.** Flow cytometry plots of KLRG-1 and CD127 expression on adoptively transferred CD-10, CD-24 and ND OT-I T_EFF_ on day 7 p.i. **G.** Percentage of MPEC and SLEC. **H.** Percentage of mAb-stimulated OT-I T_EFF_ in spleens on day 7 p.i. with LCMV-OVA. Data in A, B, D, G, and H are pooled from at least two independent experiments and analyzed using ANOVA with Dunnett’s post-test (B, G) and Student’s *t*-test (A, D, H). *, p < 0.05; **, p < 0.01; ***, p < 0.001.

**Figure S6.**
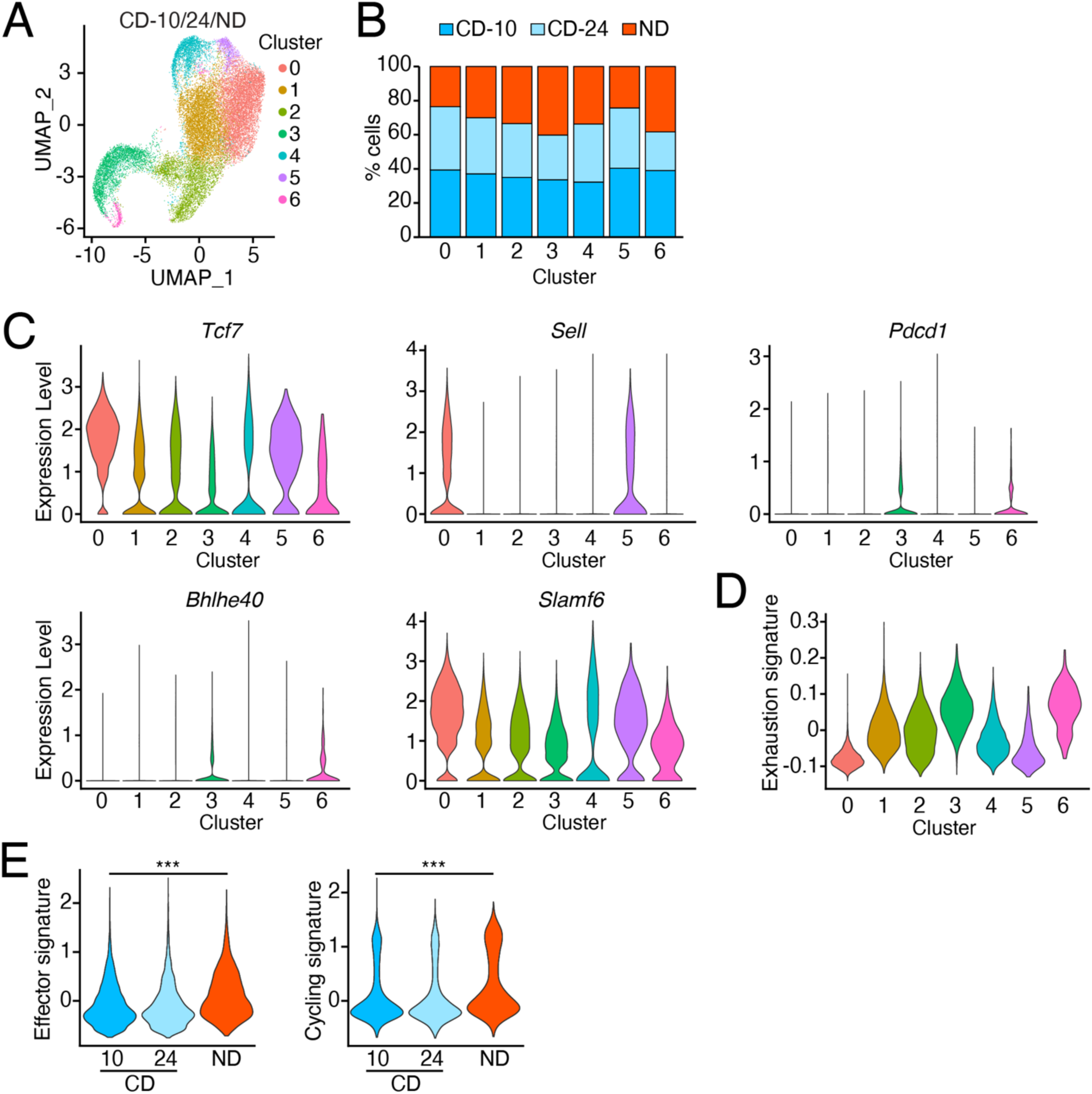
Gene expression profile of T_EFF_. **A.** UMAP projection of 24,440 sequenced CD-10, CD-24 and ND OT-I T_EFF_ isolated from spleens on day 7 p.i. Each dot corresponds to a single cell, colored according to clusters identified in an unbiased fashion using the Louvain algorithm. **B.** Cluster composition expressed as the percentage of CD-10, CD-24 and ND OT-I T_EFF_ populating each cluster. **C.** Expression level of selected genes in clusters 0-6 pooled from scRNAseq data of adoptively transferred CD-10, CD-24 and ND OT-I T cells isolated at day 7 p.i. with LCMV-OVA. **D.** Violin plots showing the exhaustion signature expression per cluster. **E.** Effector and cycling signature in CD-10, CD-24, ND OT-I T_EFF_ at day 7 p.i. with LCMV-OVA and analyzed using ANOVA with Dunnett’s post-test. ***, p < 0.001.

## Supplemental Videos

**Video S1. Intravital imaging of OVA_257-264_-pulsed DC interactions with OT-I T cells transferred at 4-8 and 22-28 h before recording.** Arrowheads depict interacting T cells. Time in min:s.

**Video S2. Under agarose imaging of control, 4 and 24 h-interacting OT-I - OVA_257-264_-pulsed DCs.** Plates were coated with CCL21 and ICAM-1. Time in min:s.

**Video S3. Under agarose imaging of Calbryte-loaded OT-I T cells and OVA_257-264_-pulsed DCs at 4 and 24 h of interactions.** Plates were coated with CCL21 and ICAM-1. Time in min:s.

**Video S4. Microfluidic chamber imaging of DOCK2-GFP OT-I T cells and OVA_257-264_-pulsed DCs at 4 and 24 h of interactions.** Arrowheads depict detaching T cells. Time in min:s.

**Video S5. Microfluidic chamber imaging of WT and DOCK2^-/-^ OT-I - OVA_257-264_-pulsed DCs at 24 h of interactions.** Arrowheads depict detaching T cells. Time in min:s.

